# SYNCAS-Mediated CRISPR-Cas9 Genome Editing in the Jewel Wasp, *Nasonia vitripennis*

**DOI:** 10.1101/2025.03.15.643426

**Authors:** Filippo Guerra, Sander De Rouck, Eveline C. Verhulst

## Abstract

Genetic engineering is a formidable approach to study biology. The development of CRISPR-Cas9 has allowed the genetic engineering of insect species from several orders, and in some species, this tool is used routinely for genetic research. However, insect gene editing often relies on the delivery of CRISPR-Cas9 components via embryo injection. This technique has a limitation: some species lay their eggs inside hard substrates or living hosts, making embryo collection impossible or labour-intensive. Recently, a variety of techniques that exploit maternal injection of nucleases have been developed to circumvent embryo injection. Yet, despite this variety of maternal delivery techniques, some insects remain refractory to gene editing. One of these is the parasitoid wasp, *Nasonia vitripennis*, an important hymenopteran model species. In this study, a recently developed method termed SYNCAS was used to perform knock-out (KO) of the *cinnabar* gene in this wasp, obtaining KO efficiencies up to ten times higher than reported for other maternal injection approaches. We found up to 2.73% of all offspring to display a KO phenotype, and we obtained up to 68 KO offspring per 100 injected mothers. The optimal timing of injection and provision of hosts for egg laying was determined. With this protocol, routine applications of CRISPR-Cas9 become feasible in this species, allowing reverse genetics studies of genes with unknown associated phenotypes and paving the way for more advanced editing techniques.

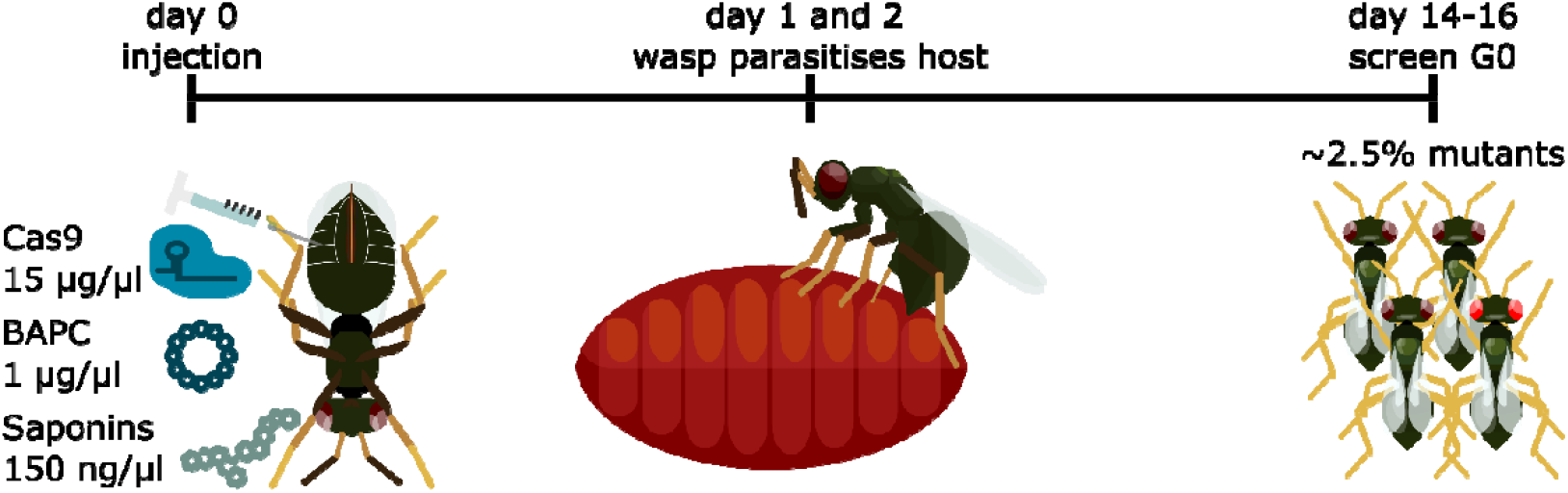

1. Maternal injection of Cas9 RNP together with BAPC and saponins (SYNCAS) results in efficient gene editing of *Nasonia vitripennis* offspring
2. Timing of injection and egg collection is important for maximising efficiency and reducing screening effort
3. The efficient genetic engineering of *Nasonia vitripennis* suggests the applicability of SYNCAS to other parasitoid wasps, a category of insects particularly difficult to genetically engineer with embryonic Cas9 injections.

## Introduction

The remarkable diversity of insects presents both an intriguing subject of study and a significant challenge for the development of standardized research methodologies. In particular, the difficulty in obtaining gene-edited insects frustrates entomologists just as much as it surprises drosophilists. Indeed, thousands of *Drosophila* lines are available, yet many species of insects remain refractory to being genetically engineered. Recent advances, including the implementation of CRISPR-Cas9-based approaches, have partially addressed these challenges. CRISPR-Cas9 relies on a Cas9 nuclease directed by a guide RNA to induce targeted DNA modifications, offering a powerful tool for genome editing. Cas9 ribonucleoprotein (RNP) complex is usually injected in early embryonic stages, before the onset of nuclear division (Burger et al., 2016). However, embryos are not always readily accessible and microinjections can cause high lethality. This is true for parasitoid species that lay their eggs inside living hosts, species that lay eggs into hard substrates, or species that lay minuscule eggs.

To circumvent these limitations, researchers developed alternative approaches that bypass direct embryonic manipulation and rely on maternal injection of CRISPR-Cas9 components to target the offspring. Some species are efficiently modified upon maternal injections of Cas9 RNP alone, a technique termed direct parental (DIPA) CRISPR. Insects as diverse as *Blattella germanica, Tribolium castaneum* (Shirai et al., 2022), *Aedes aegypti* (Shirai et al., 2023), *Sogatella furcifera* (M. Zhang et al., 2024), and *Orius strigicollis* (Matsuda et al., 2024) are efficiently modified using DIPA CRISPR. For other species, the use of Cas9 fused with a yolk-protein-derived peptide that triggers receptor-mediated uptake by the ovaries works best. This technique, termed Receptor-Mediated Ovary Transduction of Cargo (ReMOT Control), was proven efficient in the modification of mosquitoes (Chaverra-Rodriguez et al., 2018; Macias et al., 2020), *Bombyx mori* (Yu et al., 2023), *Diaphorina citri* (Chaverra-Rodriguez et al., 2023), and *Rhodnius prolixus* (Lima et al., 2024). A drawback of this method is that the yolk-peptide sequence is, to some extent, species-specific and may necessitate the production of tailor-made Cas9 depending on the target organism (Heu et al., 2020; Sharma et al., 2022). Another variation on DIPA uses chemicals to enhance Cas9 uptake by the oocytes; branched amphipathic peptide capsules (BAPC) have been used in the psyllid *Diaphorina citri* to reduce the surface charge of the RNP complex and promote its ovarian uptake (Hunter et al., 2018).

Despite the versatility and ease of use of the techniques based on maternal injection, they are no silver bullet: in the hymenopteran model *Nasonia vitripennis*, DIPA, REMOT and BAPC-assisted Cas9 delivery were shown to work (Chaverra-Rodriguez et al., 2020; X. Zhang et al., 2024) but the obtained efficiencies are insufficient to be used in practice (**Table 1**).

**Table 1:** Efficiencies of previously published techniques for *Nasonia vitripennis* genome editing. Two methods of nuclease delivery have been used for *N. vitripennis* gene editing: embryonic microinjections and maternal injections. Embryo microinjections are laborious, require micromanipulators, and result in massive embryo lethality (Li et al., 2017; personal observations). Moreover, the high efficiency obtained by Li et al. (2017) proved difficult to replicate, as shared by many researchers at the last International *Nasonia* meeting (28 July 2023, Münster, DE). Conversely, maternal injections are easy to perform, yet the low efficiency makes screening for mutants labour-intensive. In this table, “% GM offspring” reports the percentage of offspring with a knockout phenotype over the number of offspring screened, “GM wasps per 100 injections” reports how many GM wasps can be obtained per 100 injections, i.e. 100 egg microinjections or 100 maternal injections.

| Technique name | Type of injection | % GM offspring | GM wasps per 100 injections | Reference |
| --- | --- | --- | --- | --- |
| TALEN | embryonic | unknown | NA | (Lynch, 2015) |
| CRISPR-Cas9 | embryonic | 60% | 12% | (Li et al., 2017) |
| DIPA | maternal | 0.08% | 2.2% | (X. Zhang et al., 2024) |
| REMOT | maternal | 0.6% | 7.3% | (Chaverra-Rodriguez et al., 2020) |
| BAPC | maternal | 0.8% | 13.3% | (Chaverra-Rodriguez et al., 2020) |

Several aspects of *Nasonia* biology make it a highly convenient laboratory study system: it parasitizes blow fly pupae that are easily available as maggots for fishing, it has a short biological cycle (ten days at 28 °C, 14 days at 25 °C, 21 days at 20 °C) (Werren & Loehlin, 2009), and diapause can be easily induced, creating “biological backups” (Saunders, 1966). Moreover, due to the haplodiploid nature of *N. vitripennis* sex determination, unmated females lay only eggs which develop into haploid males. This eases the screening of mutants as knock-out (KO) events always produce an effect on the phenotype, even when the mutation is recessive, and simplifies the establishment of homozygous mutated lines (Lynch, 2015).

Unfortunately, trials to genetically engineer *Nasonia* resulted in impractically low efficiencies. The first attempts relied on the injection of embryos with TALENS (alleged (Lynch, 2015)) and CRISPR-Cas9 (Li et al., 2017). Subsequent efforts exploited maternal injections with REMOT technology and the use of BAPC as adjuvants (Chaverra-Rodriguez et al., 2020), and finally, DIPA (X. Zhang et al., 2024).

Recently, a technique named SYNCAS was developed to engineer arthropods with off-the-shelf nucleases and chemicals (De Rouck et al., 2024). SYNCAS relies on the use of a surfactant (saponins) and nano-capsules (BAPC) to aid protein uptake in a poorly understood, yet efficient manner. We applied the technique to knock out *cinnabar*, a gene responsible for eye colour, in *N. vitripennis*. We optimised the timing of injection in the adult female, the timing of provision of fly pupae (hosts) for egg laying, and the concentration of saponins. With this optimized protocol, we reached an efficiency of 2,73% offspring with mutated phenotype and a maximum of 67.9 mutant wasps per 100 maternal injections, which is significantly higher than previously reported studies.

## Results

### Saponin toxicity

Previous studies reported that endosomal escape is dependent on saponin concentration (Cao et al., 2020), suggesting that saponin concentration and maternally provided Cas9 efficiency could positively correlate (Macias et al., 2020). However, this chemical is toxic to *Nasonia* (Chaverra-Rodriguez et al., 2020), and a trade-off between mother survival and gene editing efficiency must be accounted for. We injected saponins in a range between 0 and 1μg/μl to define the maximum amount of saponins that would allow 70% of the wasps to lay eggs. A dose-response curve was fitted to the data (R2=0.99) (Fig 1; source data in Supplementary Table 1). Based on these results, we estimated that injections with saponins at a concentration of 150 ng/μl would result in 70% of females laying eggs. Therefore, this concentration was used as the standard for the SYNCAS formulation in *N. vitripennis*.

**Fig 1.**
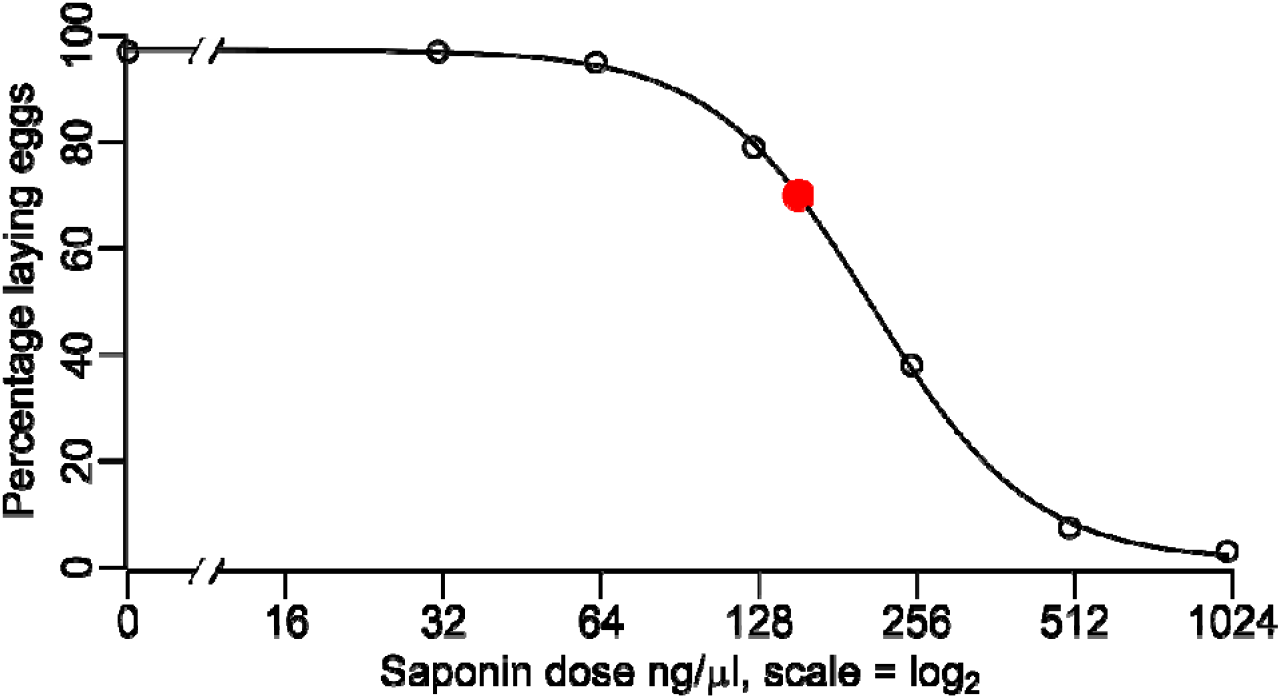
Dose-response curve for saponin effect on *N. vitripennis* egg-laying. The percentage of egg-laying females falls steeply with concentrations of saponins between 120 and 350 ng/μl. The estimated dose that would allow for 70% egg-laying ability is 150 ng/μl, and is reported as a red dot. White dots represent measured values, also reported in Supplementary Table 1. Dose-response curve and estimated dose were calculated with the R package drc (Ritz et al., 2015).

### Saponins concentration affects SYNCAS efficiency

We tested three concentrations of saponins (0 ng/μl, 150 ng/μl, and 300 ng/μl) to evaluate their impact on gene-editing efficiency, leaving the dose of Cas9, sgRNA, and BAPC constant. These formulations were injected in one-day-old virgin females. To identify the time window when most genetically engineered eggs are laid, we collected parasitised hosts three times defining three time windows: between 0 and 24 hours, between 24 and 48 hours post injection, and over 48 hours post injection. Offspring were screened for the red-eye phenotype shown in **Fig 2**. Following our previous result, saponin dose impacted survival as well as egg-laying capability and had a great impact on the production of genetically engineered offspring (**Table 2**).

**Table 2:** Effects of saponin concentration on *N. vitripennis* history traits and SYNCAS efficiency. Saponins aid the genetic engineering of *N. vitripennis* embryos, yet have a detrimental effect on the survival of injected females. A concentration of 150 ng/μl resulted in more than 2% genetically modified (GM) offspring over a 48-hour-long period and gave the best yield. All the offspring were screened for eye colour mutations. The total number of offspring (# offspring) for each treatment and time interval was determined by multiplying the average number of wasps of 12 randomly selected tubes by the total number of tubes containing offspring.

| Saponin Concentration [ng/μl] | Timepoint [hours] | Surviving females | Tubes with offspring | females producing GMO | # offspring | GM offspring | % GMO |
| --- | --- | --- | --- | --- | --- | --- | --- |
| 0 | 0-24 | 51/51 | 45/51 | 1 | 1428 | 1 | 0.07% |
| 0 | 24-48 | 50/51 | 50/51 | 1 | 2030 | 1 | 0.05% |
| 0 | 48- | 50/51 | 50/51 | 0 | 3110 | 0 | 0% |
| 150 | 0-24 | 53/56 | 32/56 | 5 | 403 | 11 | 2.73% |
| 150 | 24-48 | 53/56 | 51/56 | 11 | 1204 | 27 | 2.25% |
| 150 | 48- | 53/56 | 51/56 | 0 | 2764 | 0 | 0% |
| 300 | 0-24 | 19/45 | 12/45 | 0 | 119 | 0 | 0% |
| 300 | 24-48 | 15/45 | 11/45 | 4 | 195 | 5 | 2.56% |
| 300 | 48- | 15/45 | 13/45 | 1 | 371 | 1 | 0.27% |

**Fig 2.**
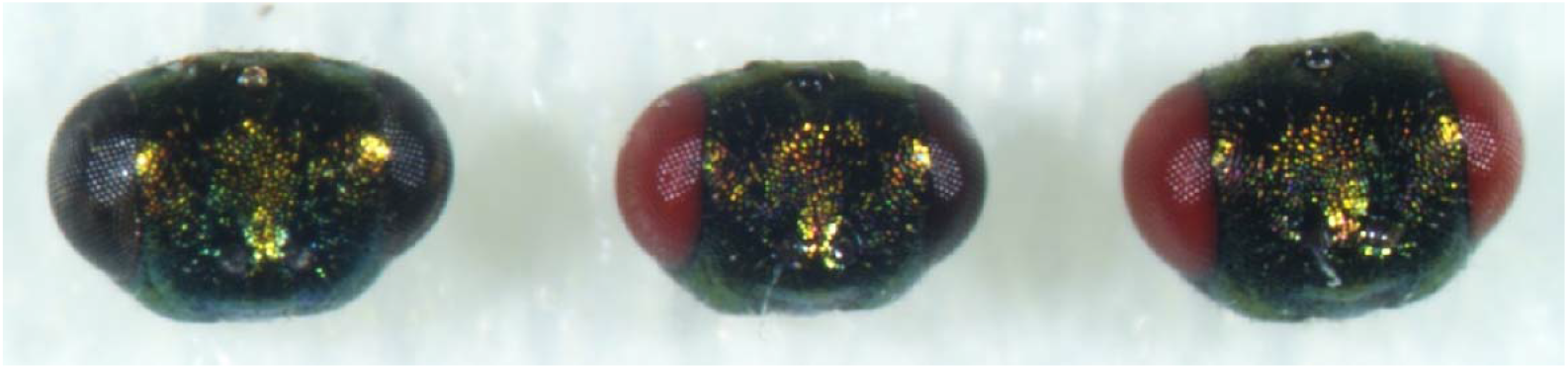
Phenotypes of G0 offspring after SYNCAS treatment. From left to right, frontal view of the head of a male with wildtype dark eyes, the head of a chimeric male having a wildtype and a red eye, and a male exhibiting the mutant phenotype, red eyes.

We calculated efficiency with two parameters: the number of mutants obtained per mother and the percentage of mutants on the total number of offspring. Injections without saponin in 51 wasps yielded two mutant males (0.04 mutants per wasp injected) of which one with chimeric phenotype, injections in 56 wasps with 150 ng/μl yielded 38 mutant males (0.68 mutants per wasp injected), and injections in 45 wasps with 300 ng/μl yielded six mutant males (0.13 mutants per wasp injected). When calculating efficiency in relation to the number of offspring, injections yielded a maximum of 0.07%, 2.73%, and 2.56% GM offspring, when injecting 0, 150, or 300 ng/μl saponins, respectively. Additionally, we identified the first 48 hours post-injection as the time window when most genetically engineered eggs are laid, highlighting the importance of timing in increasing the efficiency of the technique. This raised the question of whether injecting wasps at a different age or timing of host provision would affect efficiency.

### Injection and host provision timing both affect SYNCAS efficiency

We performed two additional injection rounds using the optimal saponins concentration of 150 ng/μl. One injection was performed in four-day-old adult females, and the second was performed in one-day-old adult females (as in the previous round of injections), but with host provision being delayed by 24 hours. Genetically engineered offspring were produced in high numbers in all conditions, yet neither per-offspring nor per-mother injected efficiency was improved (Table 3). Indeed, injections in 90 four-day-old wasps yielded 52 mutant males (0.58 mutants per wasp injected), while injections in 55 one-day-old wasps and postponing host provision yielded 27 mutant males (0.49 mutants per wasp injected). Injections in four-day-old wasps gave the biggest proportion of GM offspring between 24 and 48 hours post-injection. Injections in one-day-old wasps and postponing host provision resulted in the highest proportion of genetically engineered offspring between 48- and 72 hours post-injection.

**Table 3:** The effect of injection and host provision timing on SYNCAS efficiency. Changing the age of the wasp at injection time and postponing host provision did not improve the efficiency compared to the first round of injections, reported here again for ease of comparison. The total number of offspring (# offspring) was determined by multiplying the average number of wasps of 12 randomly selected tubes by the total number of tubes containing offspring.

| Wasp age [days] | Timepoint [hours] | Surviving females | Tubes with offspring | females producing GMO | # offspring | GM offspring | % GMO |
| --- | --- | --- | --- | --- | --- | --- | --- |
| One | 0-24 | 53/56 | 32/56 | 5 | 403 | 11 | 2.73% |
| One | 24-48 | 53/56 | 51/56 | 11 | 1204 | 27 | 2.25% |
| One | 48- | 53/56 | 51/56 | 0 | 2764 | 0 | 0% |
| Four | 0-24 | 72/90 | 77/90 | 2 | 2895 | 3 | 0.10% |
| Four | 24-48 | 72/90 | 72/90 | 7 | 1915 | 18 | 0.94% |
| Four | 48- | 72/90 | 72/90 | 11 | 3780 | 31 | 0.82% |
| One, delayed rehosting | 24-48 | 50/55 | 41/55 | 5 | 778 | 6 | 0.77% |
| One, delayed rehosting | 48-72 | 47/55 | 43/55 | 6 | 1123 | 16 | 1.42% |
| One, delayed rehosting | 72- | 46/55 | 44/55 | 1 | 2174 | 5 | 0.23% |

### Chimerism and mutation heritability

We defined the heritability of the mutation by mating the G0, red-eye males with wild-type females, and allowing the resulting G1 females to lay male G2 offspring. The presence of red-eye G2 would therefore confirm the transmission of the mutation. The results, reported in **Table 4**, suggest that editing resulting from the use of saponins is transmitted at a higher rate compared to those obtained by injections of Cas9 and BAPC alone, although the reduced sample size doesn’t allow for a definitive answer. Indeed, mutant wasps obtained from injections without saponins sired no mutant offspring, whereas mutants obtained from SYNCAS injections transmitted the mutation at rates between 83 and 96 %. Different SYNCAS treatments (i.e. injections with saponins) do not differ in mutation transmission rates [*X*^2^ (4, *N*=104)=1.5204, *p* > 0.5].

**Table 4:**
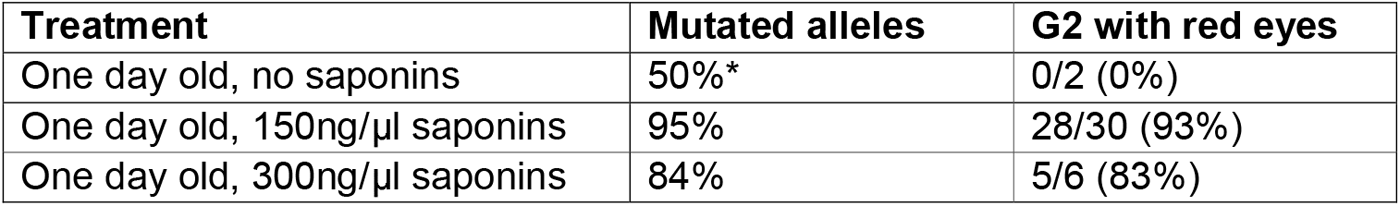
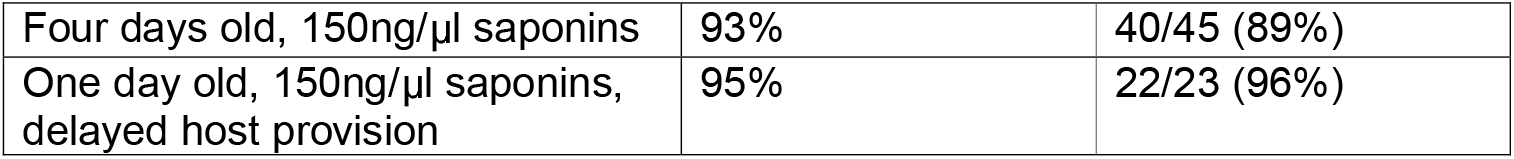
Percentage of mutated alleles and G2 offspring in different SYNCAS treatments. The percentage of mutated alleles was inferred from Nanopore sequencing, except for the treatment “One day old, no saponins”, where amplicons were individually sequenced with Sanger technology (*), revealing the chimeric nature of both individuals. The transmission of the mutation was inferred by the presence of G2 offspring with red eyes.

To define the percentage of mutated alleles in G0 red-eye males, we amplified via PCR a fragment of *cinnabar* encompassing the Cas9 target site, then we pooled the amplicons per treatment and mass-sequenced them using Nanopore technology.

Injections with no saponins resulted in two chimeric offspring and 50% of all alleles being mutated. Sequencing showed that 83 to 95% of the amplicons obtained from red-eyed males deriving from injections with saponins were mutated (**Table 4**), and had in-dels between +10 and -11 bp in size (**Supplementary Fig. 1**). As expected, the transmission of the mutation positively correlates with the proportion of mutated alleles detected in the mass sequencing.

## Discussion

Previously tested techniques relying on maternal injections achieved in *N. vitripennis* a maximum efficiency of 0.8% genetically engineered offspring, or 13.3 genetically engineered wasps per 100 injections (Chaverra-Rodriguez et al., 2020). Applying SYNCAS increased efficiency to 2.7% of offspring being genetically engineered, totalling 68 mutant wasps every 100 injections. Importantly, around one-third of all injected females laid at least one genetically engineered embryo, suggesting that injecting a small number of wasps should be sufficient to obtain at least one mutant. For example, by injecting ten females, the probability of finding at least one genetically engineered son should be above 98%.

SYNCAS is based on the synergistic action of saponins and BAPC (De Rouck et al., 2024). Although it is versatile and applicable to many arthropod species, saponin toxicity requires optimisation of the formulation composition. Indeed, Chaverra-Rodriguez and colleagues (2020) reported detrimental effects on *N. vitripennis* survival, egg-laying, and offspring viability at saponin concentrations as low as 1 ng/μl, with only 20% of the females being alive three days after injection. In our experience, these negative effects have similar severity at higher concentrations (between 250 and 500 ng/μl), and other secondary effects reported by Chaverra-Rodriguez and colleagues, such as induction of diapause in the offspring, are completely missing. Saponins are required to obtain efficient gene editing as the addition of saponins at 150 ng/μl increases efficiency more than 34-fold (0.08% vs 2.73%). Higher concentrations of saponins (300 ng/μl) did not increase the gene editing efficiency, yet it caused higher mortality and consequently less offspring. Therefore, we speculate that the maximum efficiency for this SYNCAS formulation has been reached, and lower concentrations of saponins might result in similar efficiency and cause lower lethality, reducing the number of injections necessary to generate a mutant.

We did not optimise the concentration of the other SYNCAS components (RNP and BAPC), although our previous experience suggests that high RNP concentrations are necessary to achieve high efficiency (De Rouck et al., 2024). Nonetheless, other researchers found that reducing the concentration of RNP from 4 to 2 μg/μl when performing DIPA did not result in an efficiency change (X. Zhang et al., 2024). Moreover, the efficiency they reported is similar to the one we obtained with our 0 ng/μl saponin treatment, even though we used a concentration of Cas9 substantially higher (15 μg/μl). Therefore, we suggest experimenting with lowering the concentration of RNP when using our SYNCAS formulation.

The efficiency we reported for the mutation of c*innabar* should apply to other *N. vitripennis* target genes. Yet, for genes that do not have a known/visible phenotype, screening might remain hard even with efficiencies above 2.5%. To ease this task, one might consider co-injecting Cas9-RNP against a visible marker (e.g. *cinnabar*) and the desired target, as was already done for other species modified with SYNCAS (De Rouck et al., 2024). Xinmi Zhang and collaborators (2024) found that simultaneously injecting *N. vitripennis* mothers with two sgRNAs targeting sites at a few hundred base pair distance often caused a large deletion, suggesting that two distinct cutting events happened. Moreover, the genetically engineered offspring are not uniformly distributed among the hosts, (**Supplementary Fig. 2**), and egg clutches with one genetically engineered egg are likely to contain additional ones. Thus, screening for mutants of the gene of interest only in broods having a mutant for a visible marker would alleviate the workload. In conclusion, we have demonstrated the efficiency of SYNCAS in the genetic engineering of yet another order of arthropods where embryo injections are cumbersome, and we urge scientists working on other parasitoid species to exploit this powerful technique.

### Experimental Procedures

We conducted experiments to examine the impact of two key factors on SYNCAS efficiency: the concentration of saponins and the time after oviposition when a female lays the highest number of genetically engineered eggs. An overview of the experimental procedure is provided in **Supplementary Fig. 3**.

### Insect strains and culturing

*Nasonia vitripennis* (strain AsymCx) wasps were held at 25 °C and provided daily for 4 hours with two hosts (*Calliphora vomitoria* pupae) to synchronise the age of the offspring. Ten days after oviposition, female *N. vitripennis* pupae were collected from the hosts and placed in a tube until eclosion. This was done to ensure that the females were virgins and capable of laying only haploid eggs. The wasps that eclosed within an 8-hour window were transferred to another tube and were provided with 50% honey in water and fresh hosts to allow feeding. If not otherwise specified, 24 hours later the wasps were injected with treatment solutions.

### General aspects of injection

Injections were performed under a stereomicroscope, using a capillary pulled needle (borosilicate glass capillaries, 3.5” Drummond #3-000-203-G/X), and a Femtojet 4i (Eppendorf, 5252000021). Females were anaesthetised on a CO_2_ pad and injected in the abdomen, ventral side, between sternites 3 and 4. Each wasp received around 200 nl of solution. The wasps were immediately provided with one host. Females were then moved to an incubator at 25 °C and 16/8 hours light/dark cycle. The process was performed by two people, allowing for the injection of ∼90 wasps per hour.

### Saponin toxicity

To test saponin toxicity, we injected solutions with 0, 31, 61, 125, 250, 500 and 1000 ng/µl of saponin (Merck, SAE0073) in PBS (Oxoid, BR0014G). Around 30 females per treatment were housed and injected as previously described and provided immediately with three hosts and placed in an incubator at 25 °C and 16/8 hours light/dark cycle. The survival rate was checked at 24 and 48 hours post-injection, while the parasitisation rate was calculated after 20 days.

### SYNCAS mixture preparation

After the establishment of saponin toxicity, we tested whether saponins and BAPC would increase the gene editing efficiency of Cas9 in *N. vitripennis*.

Recombinant *Streptococcus pyogenes* Cas9 protein (Alt-R® S.p. Cas9 Nuclease V3) was purchased from Integrated DNA Technologies (Leuven, Belgium) at a custom concentration of 50 μg/μl.

A sgRNA was used with protospacer sequence 5’-AACCCATCTGAACAAGCTCC-3’, specific to the eye-pigmentation gene *cinnabar*, as it was previously demonstrated to work in vivo (Li et al., 2017). Single guide RNAs (sgRNAs) were ordered from IDT (ALT-R CRISPR Cas9 sgRNA, 10 nmol). The sgRNAs were dissolved in nuclease-free water to a final concentration of 10 μg/μl.

CRISPR/Cas9 Ribonucleoprotein particles (RNPs) were prepared by mixing 3 μl of 50 μg/μl Cas9 nuclease with 5 μl of 10 μg/μl sgRNA. The mixture was incubated for 10 min at room temperature. After incubation, 1 μl of BAPC at 10 μg/μl (Phoreus Biotech), and 1 μl of saponins (either 0, 1.5 or 3 µg/μl) were added, in this order. A 30-minute incubation on ice followed. Finally, the injection mix was centrifuged at 4 °C for 10 min at 20,000 g and kept on ice until use. The final concentrations of each component in the different mixtures used were 15 μg/μl Cas9, 5 μg/μl sgRNA, 1 μg/μl BAPC, and a variable concentration of saponins (0, 150, or 300 ng/μl).

### Injections of SYNCAS mixtures

Between 45 and 90 virgin females were injected per treatment; each wasp was injected with ∼200 nl of solution. Except when otherwise specified, the wasps were immediately placed in single tubes and provided with a host. 24 hours later the hosts were collected in individual tubes labelled with a female-specific code and a second host was provided to the surviving females. 24 hours later, the hosts were similarly collected and surviving females were provided with three hosts. They were not re-hosted after this point. This setup allowed us to define the optimal time point to screen for genetically engineered offspring.

### Offspring screening for gene editing events

All G0 red-eyed males were collected in single tubes labelled with information regarding the mother’s ID and time window of egg laying. Before being euthanised for DNA extraction and testing, red-eyed males were put in individual tubes with three virgin females, which were then provided with 3-5 hosts. Ten days later, five G1 female pupae per mating were put in a single tube and upon emergence were provided with hosts. The resulting all-male G2 was screened for the presence of red-eye offspring as that would confirm the heritability of the mutation.

DNA was extracted by placing G0 red-eye males in PCR tubes with 200 μl of freshly prepared 1.25% NH_4_OH (following Wielinga et al., 2006). Tubes were heated at 99°C for 20 min, briefly centrifuged (10–30 s), then reheated with open lids at 90°C until ∼150 μl remained (∼20 min). After another brief centrifugation, samples were stored at 4 °C. The procedure was conducted in a fume hood. Each sample was used in *cinnabar* PCR amplification (M0492S, NEB Q5 polymerase) using forward (CTCTACGAGTACCGCTCAGG) and reverse (TGCAAGATCGAATCGTAACGC) primers, yielding ∼520 bp amplicons. The amplicons were loaded on a 1% agarose gel to confirm specific and equal amplification and were then pooled per SYNCAS treatment and purified (28104, QIAGEN QIAquick PCR Purification Kit). Finally, amplicons were mass-sequenced using Nanopore sequencing (Premium PCR sequencing, Plasmidsaurus, USA). Sequence alignments and KO allele frequencies were then analysed using the dedicated tools in Geneious Prime 2025.0.3.

## Acknowledgements

We would like to thank Rick Dekker who helped with collecting preliminary data on saponin toxicity and helped perform SYNCAS injections and wasp phenotyping. We also thank Prof. Thomas Van Leeuwen, Ghent University, for donating the Cas9 protein used in the experiments.

## Supplementary

**Supplementary Table 1:** Effects of saponin injections on *N. vitripennis* survival and oviposition ability. A twofold dilution series was tested for effects on the survival and oviposition rate of injected wasps. The estimated dose that would result in the desired oviposition rate of 70% is around 150 ng/μl. h P.I. = hours post-injection.

| Saponin [ng/μl] | # Wasps injected | Alive 24h P.I. | Alive 48h P.I. | Ovipositing |
| --- | --- | --- | --- | --- |
| 0 | 34 | 33 | 33 | 33 (97%) |
| 31 | 33 | 33 | 32 | 32 (97%) |
| 63 | 20 | 20 | 19 | 19 (95%) |
| 125 | 29 | 28 | 21 | 23 (79%) |
| 250 | 34 | 14 | 13 | 13 (38%) |
| 500 | 27 | 5 | 3 | 2 (7.4%) |
| 1000 | 33 | 2 | 0 | 1 (3%) |

**Supplementary Table 2:** SYNCAS formulations and injection time effects on mutation transmission rate. Different treatments result in *cinnabar* mutants that can transmit the mutation to the next generations: in particular, they do not differ in mutation transmission rate, suggesting the level of chimerism is similar between treatments. The *chi*-square statistic is 1.5204. The *p*-value is > 0.67. The differences between groups do not deviate significantly from what could be expected by chance alone.

| Chi-square calculation for mutation transmission rate |  |  |  |
| --- | --- | --- | --- |
|  | Have GM offspring | No GM offspring | Row Totals |
| Day 1, 150 ng/μl sap | 28 (27.40) [0.01] | 2 (2.60) [0.14] | 30 |
| Day 1, 300 ng/μl sap | 5 (5.48) [0.04] | 1 (0.52) [0.45] | 6 |
| Day 4, 150 ng/μl | 40 (41.11) [0.03] | 5 (3.89) [0.31] | 45 |
| Day 1, delayed ,150 ng/μl | 22 (21.01) [0.05] | 1 (1.99) [0.49] | 23 |
| Column Totals | 95 | 9 | 104 (Grand Total) |

**Supplementary Fig. 1:**
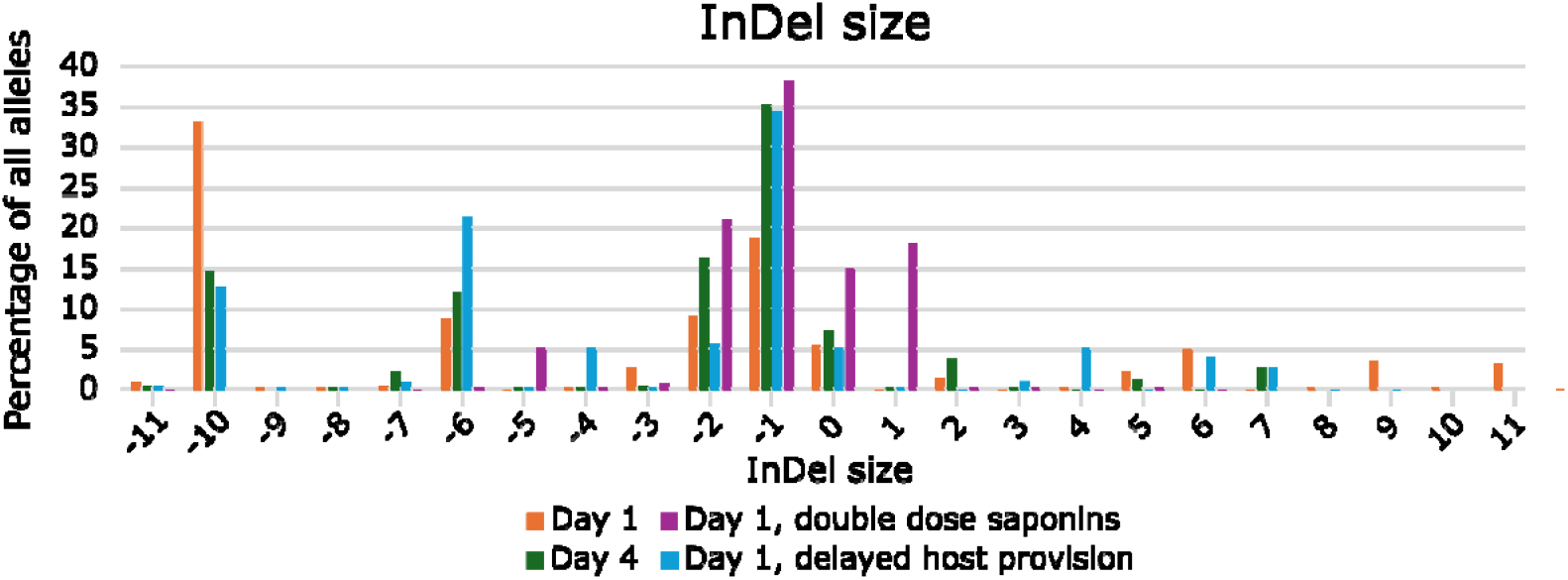
Size of insertions and deletions (InDels) around the Cas9 cutting site. The InDel size is dependent on the DNA repair mechanism and multiple mutations are generated with indel size between -10 and +11 base pairs. In orange, is the analysis of mutant offspring of one-day-old wasps injected with SYNCAS at 150 ng/μl saponins (4781 reads from 38 mutants); in green is the analysis of four-day-old wasps injected with SYNCAS at 150 ng/μl saponins (4549 reads from 52 mutants); in purple is the analysis of one day old injected with SYNCAS at 300 ng/μl saponins (3784 reads from 6 mutants); in blue is the analysis of one day old injected with SYNCAS at 150 ng/μl saponins and delaying host provision by 24 hours (4684 reads from 37 mutants). Graph was generated with Microsoft Excel (https://office.microsoft.com/excel).

**Supplementary Fig. 2:**
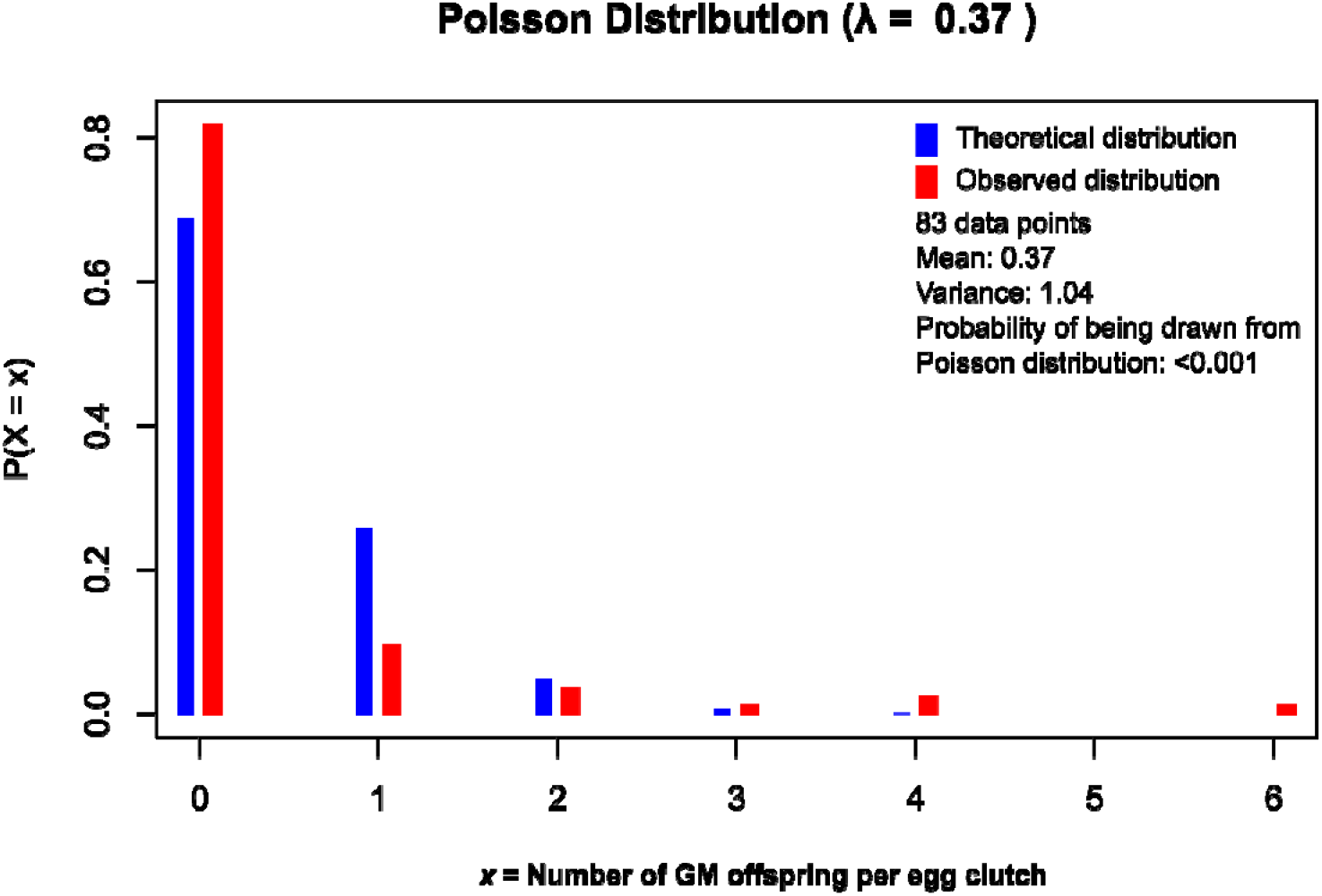
Distribution of genetically modified offspring number in parasitized hosts. When screening for a mutation, the probability of finding an egg clutch with at least one GM egg is 18%. The probability of finding a second mutated wasp in an egg clutch with a GM wasp that has at least one wasp with a mutant marker (e.g., red eyes), is 47%. This suggests that the identification of egg clutches containing GM wasps by co-injecting Cas9 against a visible marker and a second gene of interest that does not have an associated phenotype would aid the screening and identification of these mutants. The data shown in the graph are relative to the egg clutches from injections in one-day-old females with 150 ng/ μl saponins and collected between 0 and 48 hours post-injection. Image generated with R v4.4.3 (R Core Team, 2021).

**Supplementary Figure 3:**
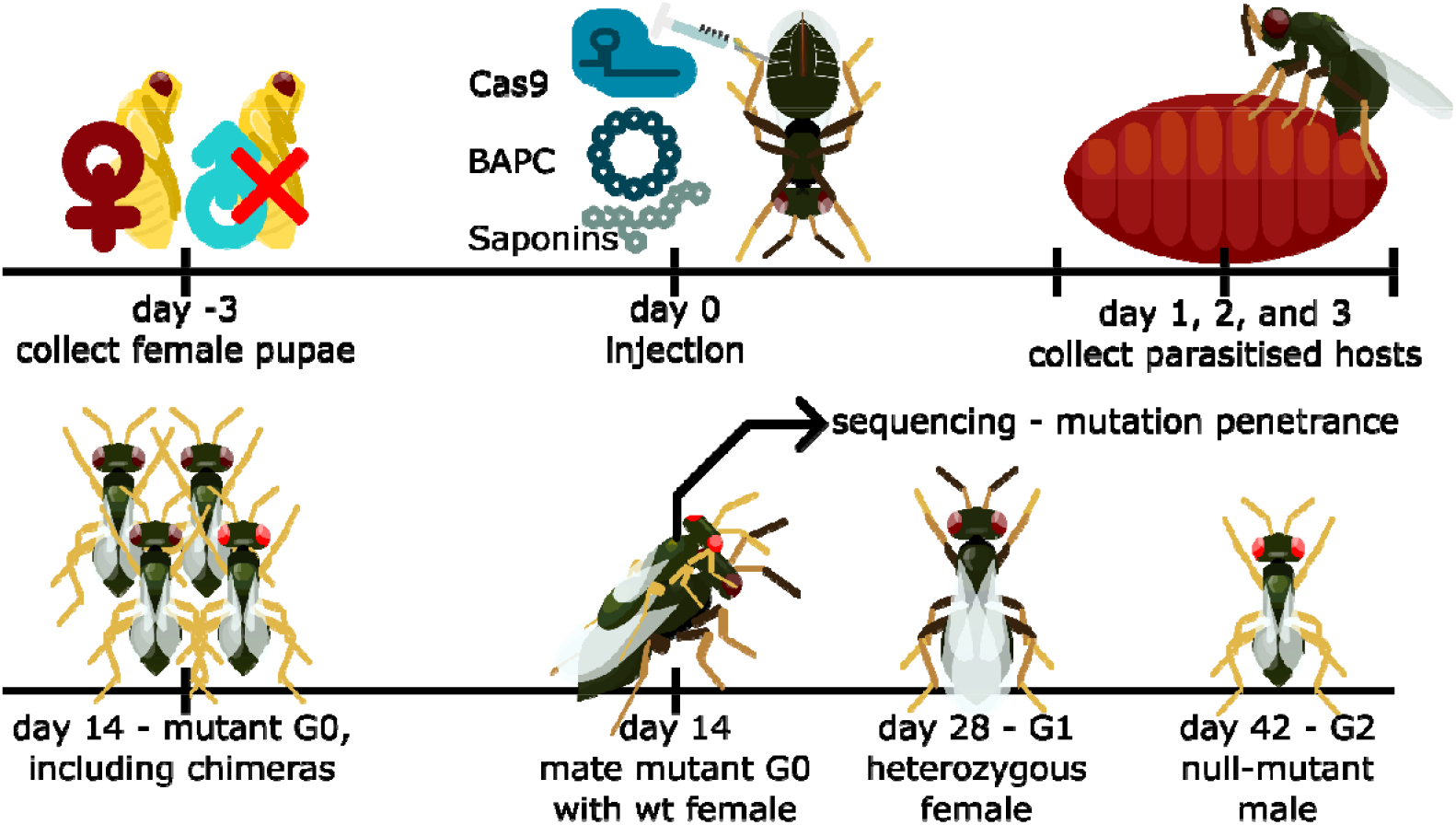
An overview of the methodology to obtain *Nasonia vitripennis* mutants using SYNCAS technology. Three days before injection, female *N. vitripennis* pupae of the same age are collected to obtain synchronized emergence of adults and ensure they are not mated and therefore capable of laying only unfertilized haploid eggs that develop into males. Wasps are injected between the third and the fourth abdominal segment with a mixture of Cas9 ribonucleoparticle, BAPC, and saponins. They are immediately provided with a host (blowfly pupae), which is collected and replaced every 24 hours. Incubation at 25 °C allows the G0 male offspring to emerge in two weeks. Screening for mutants is done by visual inspection; the progeny might contain phenotypically mutant chimaeras. To test for the heritability of the mutation and obtain null-mutant lines, the mutant males are mated with wild-type females for 24 hours. This possibly yields heterozygous G1 females, which lay null-mutant and wild-type eggs. By mating the heterozygous females with their son it is possible to obtain homozygous females and stable mutant lines. Genotypic testing via PCR amplification and sequencing of G0 informs on the penetrance of the mutation. Image generated with Inkscape v1.4 (https://inkscape.org/).

